# A novel and potent thrombolytic fusion protein consisting of anti-insoluble fibrin antibody and mutated urokinase

**DOI:** 10.1101/2020.09.06.284596

**Authors:** Shingo Hanaoka, Shinji Saijou, Yasuhiro Matsumura

## Abstract

Because the risk of thromboembolism increases with age, as well as due to infectious diseases, safer and more effective thrombolytic agents are in greater demand. Tissue plasminogen activator (tPA) is currently used clinically because it has higher binding specificity for insoluble fibrin (IF) than urokinase (UK), but even pro-tPA has catalytic activity in places other than IF. Meanwhile, UK has the advantage that it is specifically activated on IF, but it only binds IF weakly. Unlike the anti-IF monoclonal antibody (mAb) established in the past, our anti-IF mAb recognizes a pit structure formed only in IF. Here, we developed a new mAb against the pit, 1101, that does not affect coagulation or fibrinolysis, and prepared a fusion protein of UK with humanized 1101 Fab to transport UK selectively to IF. In IF-containing lesions, UK is cleaved by plasmin at two sites, Lys158/Ile159 and Lys135/Lys136. Cleavage of the former leads to activation of UK; however, because activated UK is linked by S-S bonds before and after cleavage, it is not released from the fusion. Cleavage at the latter site causes UK to leave the fusion protein; hence, we mutated Lys135/Lys136 to Gly135/Gly136 to prevent release of UK. This engineered UK-antibody fusion, AMU1114, significantly decreased the systemic side effects of UK *in vivo*. In a mouse thrombus formation experiment, the vascular patency rate was 0% (0/10) in the control, 50% (5/10) in the tPA, and 90% (9/10) in the AMU1114 treatment group. These data support future clinical development of AMU1114.

## Introduction

Hypercoagulation occurs not only in cardiovascular diseases, but also in cancer and severe infectious diseases such as influenza and coronavirus infection, worsening their pathologies.^1-3^ In patients with such severe conditions, administration of thrombolytic agents should be carried out with caution, and safer forms of administration are desirable. Currently, tissue plasminogen activator (tPA) is the thrombolytic agent used most commonly in clinics around the world because it binds IF more specifically than UK.^4^ However, even under tPA treatment, bleeding is a clinically serious side effect.^5^ To address this issue, efforts have been made to increase the fibrinolytic activity of plasminogen activators by selectively targeting them to IF in lesions. For these purposes, a mAb against IF called 59D8 is utilized as a delivery tool.^6^ For example, some groups prepared a chemical conjugate of pro-UK with this mAb.^7,8^ Another group produced a recombinant fusion protein of the catalytic domain of UK and the scFv of 59D8.^9^ Still other groups developed chemical conjugates of tPA and 59D8.^10,11^

However, for several reasons including low yield, inconsistent coupling, and low superiority relative to the original plasminogen activators, none of the fusions or conjugates were evaluated clinically. In addition to technical problems related to chemical conjugation and protein fusion, the 59D8 mAb used in those groups bound not only IF but also fibrinogen, which circulates in the blood. Therefore, it is assumed that those formulations were not efficiently delivered to the lesion because they were neutralized by the large amounts of fibrinogen in the blood.

As part of our research into cancer and blood coagulation, we established a mAb (102-10) that recognizes only IF and not fibrinogen, soluble fibrin (fibrin monomer), or soluble fibrin degradation products (FDP).^12,13^ The epitope of 102-10 is on the β chain that underlies the pit structure formed by the hydrophobic regions of the β and γ chains, which is exposed only when IF is formed. Because fibrinogen, the fibrin monomer, and FDPs are soluble, the pit structure is completely closed in aqueous solution due to hydrophobic bonding; consequently, 102-10 does not bind soluble molecules such as fibrinogen, the fibrin monomer, or FDP. The amino acid sequence of this epitope is widely conserved among animals ranging from fish to humans. In other words, even though 102-10 is an anti-human IF antibody produced in mice, it also recognizes mice IF. This suggests that data from mouse experiments can be extrapolated to humans.

Even single-chain tPA (pro-tPA) has enzymatic activity that converts plasminogen in circulating blood into plasmin.^14,15^ Plasmin and activated tPA in the blood are inhibited by innate α2-plasmin inhibitor (α2-PI)^16^ and plasminogen activator inhibitor-1 (PAI-1),^17,18^ respectively. On the other hand, pro-UK is rarely activated naturally in blood circulation and is not inhibited by PAI-1.^19^ Consequently, UK is active only on IF in the lesion, where plasmin is abundant. Based on these observations, we hypothesized that a thrombolytic agent superior to tPA could be obtained if it were possible to efficiently deliver pro-UK to IF. Thus, we have prepared a fusion protein of pro-UK and anti-IF mAb to deliver pro-UK selectively to IF in lesions in the body.

## Methods

### Development of mAb 1101 and its humanization

The fibrinogen β-chain D domain (a.a. 228-491, UniprotKB entry number P02675) was expressed in *E. coli* and used as an immunogen. The antigen, mixed with adjuvant, was administered four times intraperitoneally to BALB/c mice, followed by final immunization through the tail vein. Three days after the final immunization, the spleen was removed, and the spleen cells were fused with X63 myeloma cells by the PEG method to obtain an antibody-producing hybridoma.

Hybridomas that were immunogen-positive, IF-positive, and fibrinogen-negative were screened by ELISA. The clone producing an antibody with the highest binding strength and specificity was established and named 1101. The isotype of the antibody was determined using an isotype-specific anti-mouse antibody measurement kit (Bethyl). An antibody that was unreactive to both IF and fibrinogen was used as a negative control. Cloning was performed by 5’-RACE, and the gene sequence was confirmed. The confirmed CDR region was inserted into human IgG (Human Monoclonal Antibodies-Methods and Protocols, Springer Protocols). Humanized 1101 and control antibodies were expressed in CHO cells, and IF specificity was confirmed by ELISA.

### Enzyme-linked immunosorbent assay (ELISA)

#### Preparation of IF-immobilized plates

Fibrinogen (Sigma-Aldrich) was immobilized on 96-well plates at a concentration of 1 μg/well and allowed to stand overnight at 4°C. One hundred microliters of 0.05 U/mL thrombin (Sigma-Aldrich), 7 mM L-cysteine, and 1 mM CaCl_2_ was added to each well, and the plates were incubated at 37°C for 2 hours. After blocking with N102 (Nichiyu) containing 10% sucrose, blocking was performed again with TBS-T (Tris-buffered saline with Tween-20) containing 1% BSA, yielding IF-immobilized plates. *ELISA*. Fibrinogen (Sigma-Aldrich), soluble fibrin (Sekisui Medical), and fibrin degradation product (Sekisui Medical), were also immobilized on 96-well plates (1 μg/well of each compound). The plate was reacted with antibody solution (1 μg/mL) for 1 hour, washed with TBS-T, reacted with HRP-conjugated anti-human IgG-Fc antibody or anti-mouse IgG-Fc antibody (Bethyl) for 1 hour, and washed with TBS-T. After incubation with the TMB (3,3’,5,5’-tetramethylbenzidine) color-developing reagent for 15 minutes, the reaction was stopped by addition of 2 N H_2_SO_4_, and absorbance at 450 nm was measured. The antibodies used in this experiment were 1101, the negative control, and the previously established 59D8 mAb, which recognizes the N-terminus of fibrinogen after thrombin cleavage.

### Fibrin gel turbidity assay

The turbidity assay was performed according to a minor modification of the method of Kim et al.^31^ at 37°C in a 96-well plate (Corning) on a SpectraMax Paradigm plate reader (Molecular Devices, Sunnyvale). Measurements were performed in quadruplicate. Turbidity was monitored once a minute at a 350-nm wavelength and calculated as the mean value (n = 4) in a volume of 100 μL HBS (HEPES [N-2-hydroxyethylpiperazine-N’-2-ethanesulfonic acid]-buffered saline solution) at pH 7.4.

For fibrin gel formation, 2.0 mg/mL fibrinogen (Sigma-Aldrich), 5 mM CaCl_2_ (Wako Pure Chemical), 0.01% Tween 80 (Sigma-Aldrich), and 0.5 NIH unit/mL thrombin (Sigma-Aldrich) were added to 10 μg/mL 1101 mAb or 3 U/mL anti-thrombin (SLS Behring K) as AT III (antithrombin III).

For the lysis of the fibrin gel, 2.0 mg/mL fibrinogen (Sigma-Aldrich), 5 mM CaCl_2_ (Wako), 0.01% Tween 80 (Sigma-Aldrich), 0.5 NIH unit/mL thrombin (Sigma-Aldrich), 0.2 μM PLG (Enzyme Research Laboratories), and 0.3 nM tPA (Alteplase, Kyowa Hakko Kirin) were added to 10 μg/mL1101 mAb or a mixture of 0.10 μM α2-PI (Hematologic Technologies) and 2.0 ng/mL PAI-1 (ProSpec), and used as a negative control.

### Immunohistochemistry of human thrombus

Human tissue sample with thrombus was generously provided by Dr. Genichiro Ishii (Pathology Division of National Cancer Center Hospital East). Immunostaining was performed on paraffin-embedded tissue. In brief, 6 μM thick sections were deparaffinized with xylene and rehydrated with decreasing concentrations of ethanol in water. Inhibition of endogenous peroxidases was performed by incubating slides in 3% hydrogen peroxide (Wako) for 20 minutes. Antigen retrieval was achieved by microwaving the slides for 10 minutes in hot (98°C) Tris-HCl buffer (TB) at pH 9.0, followed by 30 minutes of cooling down to room temperature. The sections were then washed with phosphate□buffered saline (PBS) (Wako) for 10 minutes. After blocking with 5% skimmed milk (Becton Dickinson) at room temperature for 30 minutes, the sections were incubated overnight at 4°C in a humidified chamber with 10 μg/mL of HRP-conjugated 1101 IgG antibody. After washing with PBS for 5 minutes, color development was achieved by applying diaminobenzidine tetrahydrochloride (DAB) staining reagent (Dako) for 2 minutes. The sections were then counterstained with hematoxylin (Muto Pure Chemicals), dehydrated through ethanol and xylene, and coverslipped using Mount-Quick mounting medium.

### Construction of several types of Fab-UK

UK (wild type) was linked to the Fab region of the H chain of humanized 1101 antibody via a 16-residue linker (GSGGGGSGGGGSGGSS) (Construct 1). The UK mutant (Lys135-Lys136 → Gly135-Gly136) was linked to the same Fab region to yield Construct 2 (AMU1114). The His8 tag was added to the C-termini of all constructs. Each of the prepared cDNAs was introduced into pcDNA3.3-TOPO (Invitrogen).

### Expression and purification of Fab-UK in CHO

The vectors were used for transient expression with the ExpiCHO Expression System (Thermo Fisher Scientific). FBS (20%) was added to the medium to suppress enzymatic activity. Transiently expressed culture supernatant was applied to a Superdex 75pg (GE) gel filtration column equilibrated with 50 mM Tris-HCl (pH 8.5)/300 mM NaCl. Next, the elution area of the target product was collected and applied to a Ni column using the same buffer conditions as for the Superdex 75pg. After washing with the same buffer, the target product was eluted with PBS containing 200 mM imidazole and 150 mM L-Arg. To obtain the final product, the eluted solution was purified on a Superdex 200pg (GE) gel filtration column equilibrated with PBS buffer containing 150 mM L-Arg.

### Preparation of fibrin gel

Blood was collected in a vacuum blood collection tube (Venoject II VP-P070K30, Terumo) and centrifuged in a Medifuge at room temperature for 13 minutes to collect plasma. Human fibrinogen (Sigma-Aldrich) labeled with fluorescent Alexa Fluor 647 (Thermo Fisher Scientific) was added to the collected plasma (1/100^th^ volume). Plasma mixed with AF647-labeled fibrinogen was dispensed into a 96-well plate at 140 μL per well, and then 20 NIH units/mL thrombin was added at 7.5 μL/well and left to stand at room temperature for 15 minutes. After the plasma coagulated, the sample was centrifuged at 500 *g* for 3 minutes.

### Fibrinolytic assay of Constructs 1 and 2 using the fibrin gel

UK (Novoprotein), Construct 1, and Construct 2/AMU1114 were adjusted to 0.4 μM with human plasma (Kohjin-bio), and 90 μL of the adjusted sample was added to fibrin gel. After fibrinolytic agents were added to fibrin gel labelled with Alexa Fluor 647, the mixture was reacted at 37°C for 4.5 hours, and the fluorescence eluted in the supernatant was measured at excitation and emission wavelengths of 647 and 680 nm, respectively (Ex647/Em680).

### Fibrinolytic assay of UK, tPA, and AMU1114 using fibrin gel

The molecular weights of UK (Urokinase, Mochida Pharmaceutical), tPA (Alteplase, Kyowa Hakko Kirin), and AMU1114 are 46 kDa, 59 kDa, and 96 kDa, respectively. To determine the approximate specific activity of fibrinolysis of those molecules, UK, tPA, and AMU1114 were adjusted to 1 mg/mL, 1 mg/mL, and 2 mg/mL, respectively, in PBS. Each adjusted sample was diluted 10-fold with human plasma (Kohjin-bio) and reacted at 37°C for 120 minutes. Each reacted sample was further diluted 10-fold with human plasma, and fibrin gel (150 μL/well) was added. After addition of the fibrin gel, the mixture was reacted at 37°C for 1 hours, and Alexa Fluor 647 eluted in the supernatant was measured at Ex647/Em680.

### Effects of UK, tPA, and AMU1114 on plasma level of plasminogen in mice

To observe the systemic effect of UK, tPA, and AMU1114 *in vivo*, these thrombolytic agents were administered intravenously (i.v.) in C57BL/6J mice at equal molar ratios; the *in vitro* fibrinolytic specific activities of the three agents are almost identical. According to previous evaluations of *in vitro* specific activity of fibrinolysis, UK and tPA were injected i.v. at 10 mg/kg,^20^ and AMU1114 was administered i.v. at 20 mg/kg. Each group consisted of three mice. Thirty minutes after administration, the mice were anesthetized with isoflurane, and 500 μL of blood was collected from the heart. The blood was immediately transferred to a 1.5-mL tube and cooled with a cooling agent until use. A syringe pre-filled with 50 μL of 19% sodium citrate solution was used for blood collection. The collected blood was centrifuged at 10,000 *g* for 2 minutes to collect plasma and stored at -80°C until use.

Each frozen plasma sample was thawed on ice and diluted 40-fold with TBS, and 20 μL of the diluted sample was mixed with 100 μL of 250 unit/mL UK. As a control, 20 μL of diluted sample was mixed with 100 μL TBS. The mixed sample was reacted at 37°C for 1 hour, mixed with 20 μL of substrate (Test Team SPLG, Sekisui medical), and reacted at 37°C for 6 hours. After the reaction, the absorbance at 405/505 was measured. Based on the measured absorbance, the level of plasminogen in each sample was calculated based on the level of plasminogen in plasma collected from untreated mice. All animal procedures and experiments were approved by the Committee for Animal Experimentation of the National Cancer Centre, Tokyo Japan.

### *In vivo* thrombus formation and re-dissolution of thrombi formed by thrombolytic agents

This experiment was entrusted to LSI Medience, which has extensive experience in mouse thrombosis research. Briefly, a thrombus was formed in the carotid artery of the mouse by the photo-chemically induced thrombosis (PIT) method, and each thrombolytic agent was administered to determine when the carotid artery was opened or occluded. In the PIT method, light with a wavelength of 540 nm is used to irradiate Rose Bengal, generating singlet oxygen at the irradiation site that damages endothelial cells and induces platelet aggregation, thereby promoting thrombus formation.

Ninety microliters of Alteplase (tPA) was administered to mice at 3 mg/kg body weight (average body weight, 30 g; n = 10 for each administration group); AMU1114 was administered at 90 μL at 6 mg/kg. At this dose, tPA and AMU1114 are equally active in *in vitro* systems. In each experiment, thrombus formation in the carotid artery was initiated by administration of Rose Bengal dye at 0 minutes, followed by light irradiation. Five minutes after the start of thrombus formation, PBS, tPA, or AMU1114 was administered. Irradiation was stopped 30 minutes after the start of thrombus formation, and observation was terminated 30 minutes after that. The times when the carotid artery was occluded or open were recorded over the 60-minute observation period.

### Statistical analyses

Statistical analyses of ELISA assay, fibrinolytic assay, and plasma level of plasminogen were performed using the EZR software. *P*-values were determined by analysis of variance (ANOVA) and Tukey’s test.

In animal experiments, we compared the control with tPA and control with AMU1114. We also compared tPA and AMU1114 in terms of the number of cases in which blood vessels were patent at the end of the experiment. The time to vascular occlusion and the patency rate of blood vessels were tested by the F test (significance level: 5%) for uniformity of variance. For the number of cases in which blood vessels were patent at the end of the experiment, a χ ^2^ test was performed. If the variances were uniform, Student’s t test was used for comparison. If the variances were not uniform, the comparison was performed using the Aspin-Welch t-test. Significance levels were set to 5% and 1%, and a two-sided test was performed. For the evaluation data, the average value ± standard error of each group was obtained. In addition, individuals whose blood vessels were patent at the end of the experiment were counted.

## Results

### Characterization of an anti-IF mAb

The anti-IF mAb clone 1101 was established by immunizing mice with the D domain of fibrinogen β-chain (a.a. 228-491) (Figure 1A). The mAb was humanized by analyzing the gene encoding the established antibody and inserting part of the CDR region into the gene encoding human IgG. Like 102-10, 1101 bound only IF and not soluble fibrinogen, the fibrin monomer, or FDP. On the other hand, mAb 59D8, which recognizes the N-terminus of fibrinogen cleaved by thrombin, recognized not only IF but also fibrinogen^6^ (Figure 1B). Like 102-10, 1101 bound not only human IF but also mouse IF (Figure 1C). In addition, the binding strength and specificity of 1101 for IF was higher than that of 102-10 (data not shown). Immunohistochemistry with 1101 revealed clear IF deposition in human thrombosis (Figure 1D). In the IF formation assay, anti-thrombin III (AT III) suppressed IF formation. On the other hand, 1101 caused IF formation to the same extent as control PBS. In the fibrinolysis assay system, α2-plasmin inhibitor (α2-PI) and plasminogen activator inhibitor 1 (PAI-1) clearly decreased fibrinolytic activity. As in the control, 1101 did not delay fibrinolysis. These results indicated that the coagulation and fibrinolytic systems were unaffected by 1101, as with the previously established antibody 102-10 (Figure 1E).^21^

**Figure 1.**
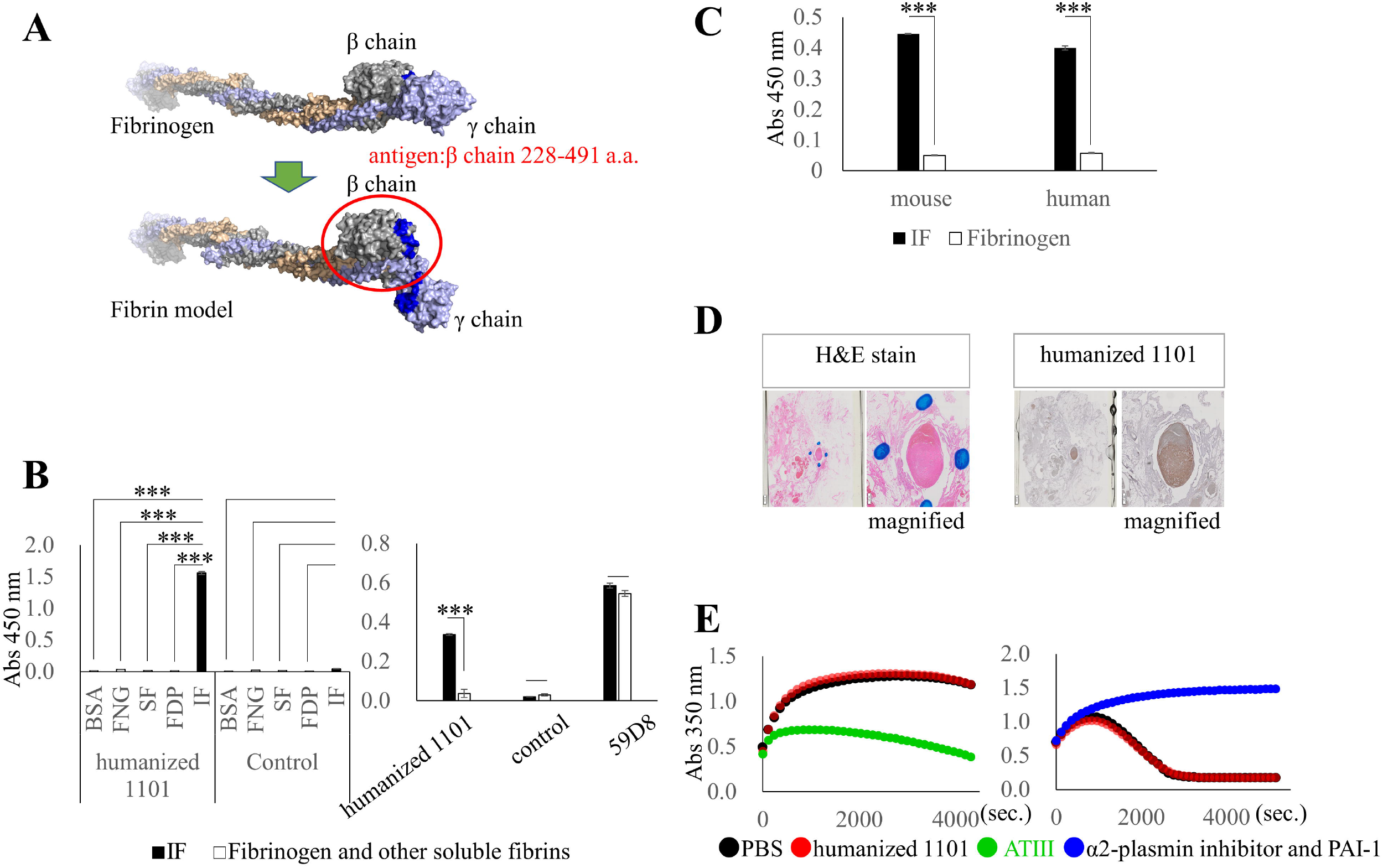
Characteristics of anti-fibrin 1101 antibody. (A) Diagram of the structural change from fibrinogen to IF. The newly identified pit structure of IF is shown in blue, and the region containing the antigen (a.a. 228-491) is circled in red. (B) ELISA (n = 3). Left: Comparison of 1101 and control antibody. BSA, bovine serum albumin (blocking agent); FNG, fibrinogen; SF, soluble fibrin; FDP, fibrin degradation product; IF, insoluble fibrin. Right: Comparison of 1101, control, and 59D8 antibodies. IF (black), fibrinogen (white). (C) ELISA (n = 4) with 1101 for human and mouse fibrinogen and IF. ELISA results are shown as means ± S.E. Statistical analysis was performed using Tukey’s test. \**P* < .05. \*\**P* < .01. \*\*\**P* < .001. (D) Immunohistochemistry of human thrombus (surrounded by blue dots) with 1101 mAb. (E) Left: Fibrin gel formation (coagulation) started by of fibrinogen, thrombin, CaCl_2_, and Tween 80 with 1101 or the coagulation inhibitor AT III. Fibrin gel in the solution was measured by monitoring turbidity (absorption at 350 nm). Right: fibrin gel degradation (fibrinolysis) was monitored in the same way as fibrin gel formation by mixing fibrinogen, thrombin, CaCl_2_, Tween 80, PLG, and tPA with 1101 or the fibrinolysis inhibitors α2-PI and PAI-1.

### Comparison of UK and its constructs

A portion of UK was linked to the Fab region of the H chain of humanized antibody 1101 via a 16-residue linker. An antibody-UK fusion protein was generated for two types of UK (Figure 2A). Construct 1 is a fusion protein of antibody and wild-type UK. Construct 2 (AMU1114) is a fusion protein of antibody and mutated UK with the Gly135/Gly136 substitutions. UK contains two sites cleaved by plasmin; one of them, Lys158/Ile159, is in the UK catalytic domain. Cleavage of this site causes activation of UK, but even if it is cleaved, active UK does not separate from the fusion protein because it is connected by an S-S bond that sandwiches the cleavage site. The other plasmin cleavage site is Lys135/Lys136, which is located on the N-terminal end of the UK catalytic domain, called the Kringle domain. Because this cleavage site is not connected by an S-S bond, cleavage by plasmin results in release of activated UK from construct 1, which would result in insufficient IF lysis. Therefore, by converting Lys135/Lys136 to Gly135/Gly136, we prevented this region of AMU1114 from being cleaved by plasmin, enabling activated UK in the fusion protein to function on IF in the lesion for a long period of time. SDS-PAGE clearly indicated a 96-kDa AMU1114 in the non-reduced condition. In the reduced condition, the 72.7-kDa heavy chain of 1101 Fab connected to the mutated UK and the 23.4-kDa light chain of 1101 Fab were obtained (Figure 2B). The mutated UK did not decrease fibrinolytic activity; the *in vitro* fibrinolytic activity of AMU1114 was similar to that of UK and construct 1 (Figure 2C).

**Figure 2.**
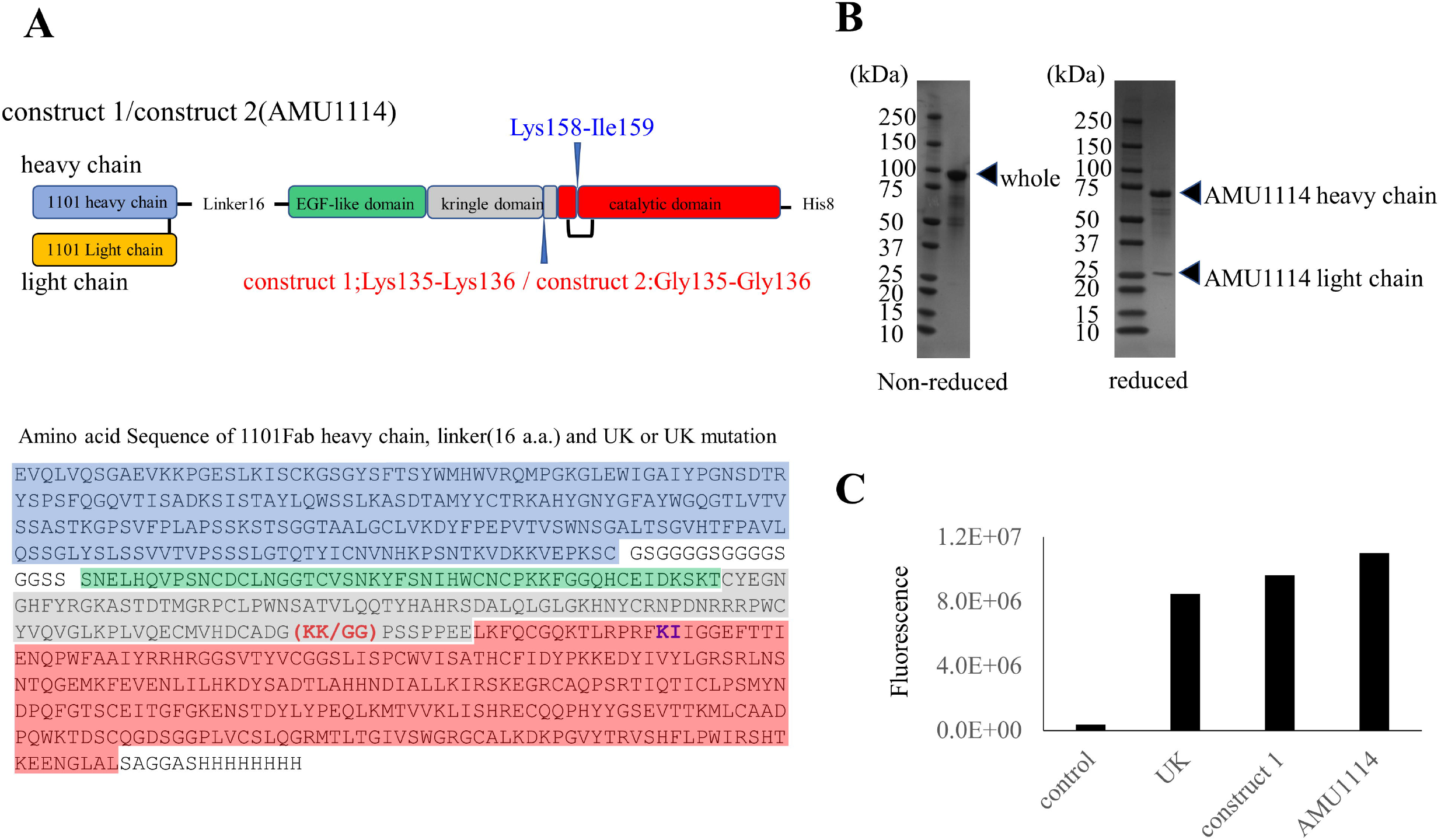
Structure of 1101 antibody-UK fusion protein. (A) Schematic diagram of two types of antibody-UK fusion constructs and amino acid sequence of the heavy chain of constructs 1 and 2. Construct 1: 1101Fab-UK; Construct 2: 1101Fab-mutated UK (AMU1114). The heavy chain of 1101 Fab is connected with the mutated UK via a 16-amino acid linker. (B) SDS-PAGE of AMU1114. Full-length (96-kDa) AMU1114 was detected in the non-reduced condition. In the reduced condition, the 72.7-kDa of heavy chain of 1101 Fab was connected with the mutated UK and the 23.4-kDa of light chain of 1101 Fab. (C) Fibrinolytic activity of UK, construct 1, and AMU1114. Fibrin dissolution analysis was performed using the fluorescent labelled fibrin gel. This experiment was conducted to confirm that mutation of UK did not change its catalytic activity.

### Fibrinolytic activity in vitro and in vivo

When comparable molar quantities of UK, tPA, and AMU1114, were applied to fibrin gels, UK, tPA, and AMU1114 all showed significantly stronger fibrinolytic activity than the control (*P* < .001), and fibrinolytic activities were similar among the three agents (*P* > .5) (Figure 3A).

**Figure 3.**
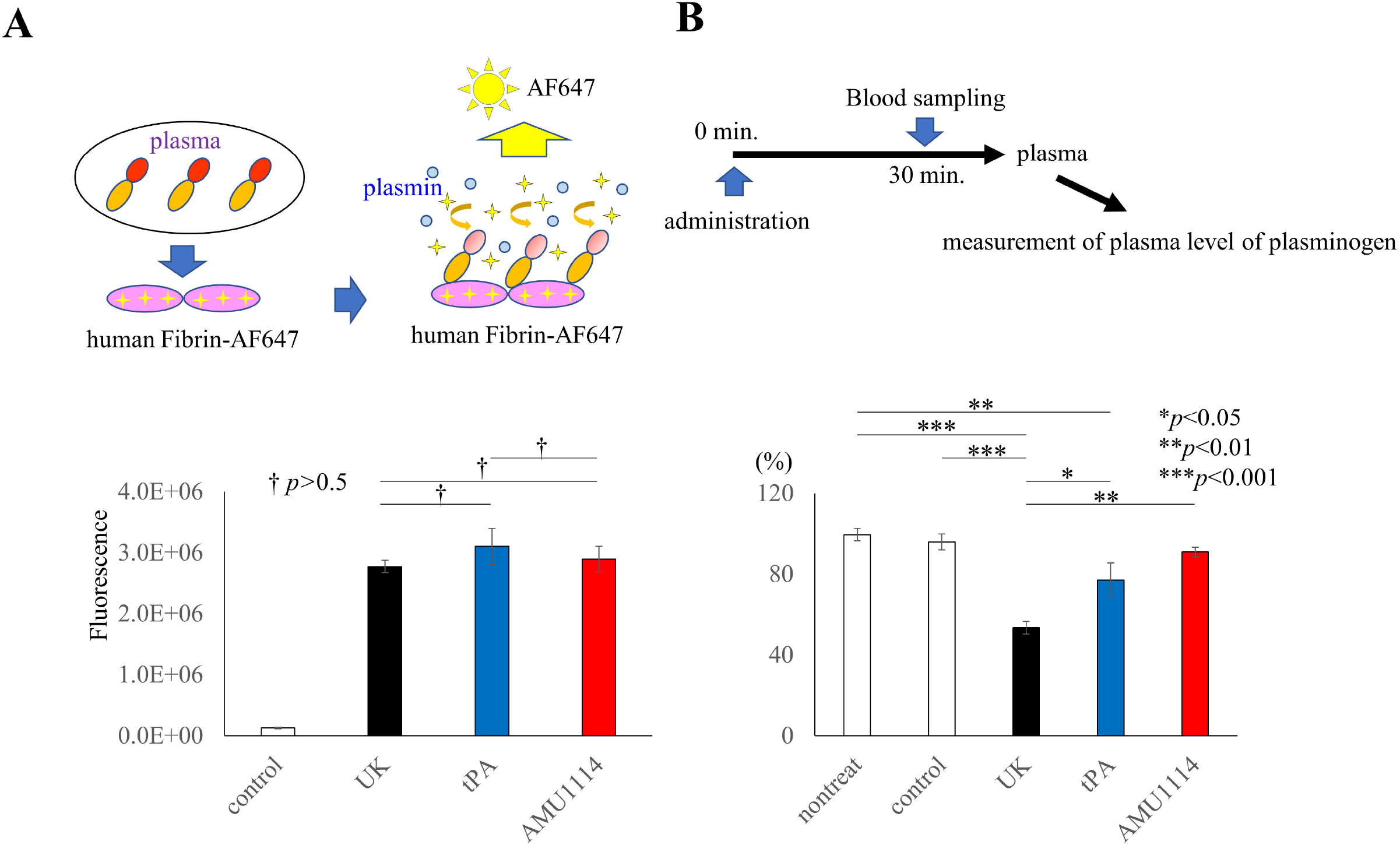
Fibrinolytic assay using fibrin gel and effect on plasma level of plasminogen in mice. (A) Fibrin dissolution analysis using fluorescently labeled fibrin gel (n = 3). Statistical analysis was performed using the Tukey’s test. (UK vs tPA, UK vs AMU1114, tPA vs AMU1114, ^†^*P* > .5) (B) Effects of various thrombolytic agents on plasma plasminogen levels in mice (n = 3). The amount of plasminogen in each plasma sample was measured, and the ratio relative to the amount of plasminogen in untreated mice is shown. Statistical analysis was performed using Tukey’s test. \**P* < .05. \*\**P* < .01. \*\*\**P* < .001.

Plasma plasminogen levels after administration of PBS, UK, tPA, and AMU1114 were 94.7%, 51.1%, 74.9%, and 85.5%, respectively, relative to the level in plasma of untreated mice. UK administration significantly decreased plasma plasminogen relative to the other groups (UK vs tPA, *P* < .05 and UK vs AMU1114, *P* < .01). There was no significant difference in plasminogen levels between tPA and AMU1114 treatment. However, plasma plasminogen was significantly lower in the tPA group than in the untreated group (*P* < .01), and there was no significant difference between the AMU1114 and untreated groups (Figure 3B).

### Comparison of antithrombotic activities

In the PIT experiment in the mouse carotid artery,^22^ the times from the start of green light irradiation to complete obstruction of blood flow in the control, tPA, and AMU1114 groups were 553.1 ± 141.9, 1135.7 ± 308.3, and 1267.8 ± 308.2 seconds, respectively. Multiple comparisons were performed among the three groups, but no significant differences were detected (Figure 4A).

**Figure 4.**
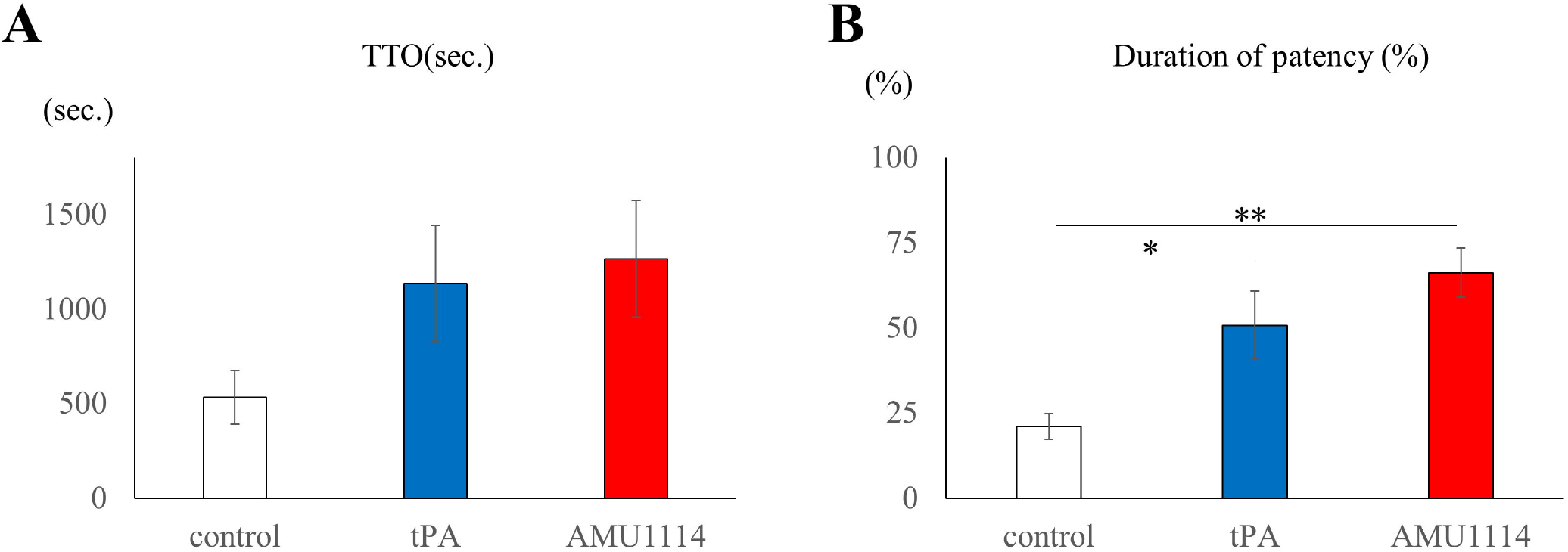
Effect of AMU1114 or Alteplase (tPA) on time to occlusion (TTO) and duration of patency. (A) Time from the start of green light irradiation to complete obstruction of blood flow in the control, tPA, and AMU114 groups. There was no significant difference between control and tPA (*P* = .0999) or between control and AMU1114 (*P* = .0501). (B) Rate of vascular patency for 60 minutes of control, tPA, and AMU1114 groups. Each value represents the mean ± S.E. *, control vs tPA *P* < .05: Significantly different from control group (Aspin-Welch t-test) **, control vs AMU1114 *P* < .01: Significantly different from control group (Student’s t-test) Not significant between tPA and AMU1114.

Rates of vascular patency for 60 minutes were 21.1 ± 3.7%, 50.9 ± 10.1%, and 66.3 ± 7.2% respectively, in the control, tPA, and AMU1114 groups. The rate of vascular patency for 60 minutes in the tPA and AMU1114 treatment groups was significantly higher than that in the control (Figure 4B). In addition, when multiple comparisons were performed among the control, tPA, and AMU1114 groups, significant differences were detected between the control and tPA group (*P* < .05) and between the control and AMU1114 group (*P* < .01), but there was no significant difference between the tPA and AMU1114 groups, although AMU1114 tended to have a higher rate of vascular patency than tPA. At the end of the measurement, the number of mice with open blood vessels was 0/10 in the control group, 5/10 in the tPA group, and 9/10 in the AMU1114 group (Figure 5A,B). Significant differences in the number of individuals whose blood vessels were patent were detected between the control and tPA groups (*P* = .0098), and between the control and AMU1114 groups (*P* = .0001). On the other hand, no significant difference was detected between the tPA and AMU1114 groups, although AMU1114 tended to have a stronger thrombolytic effect than tPA (*P* = .051).

**Figure 5.**
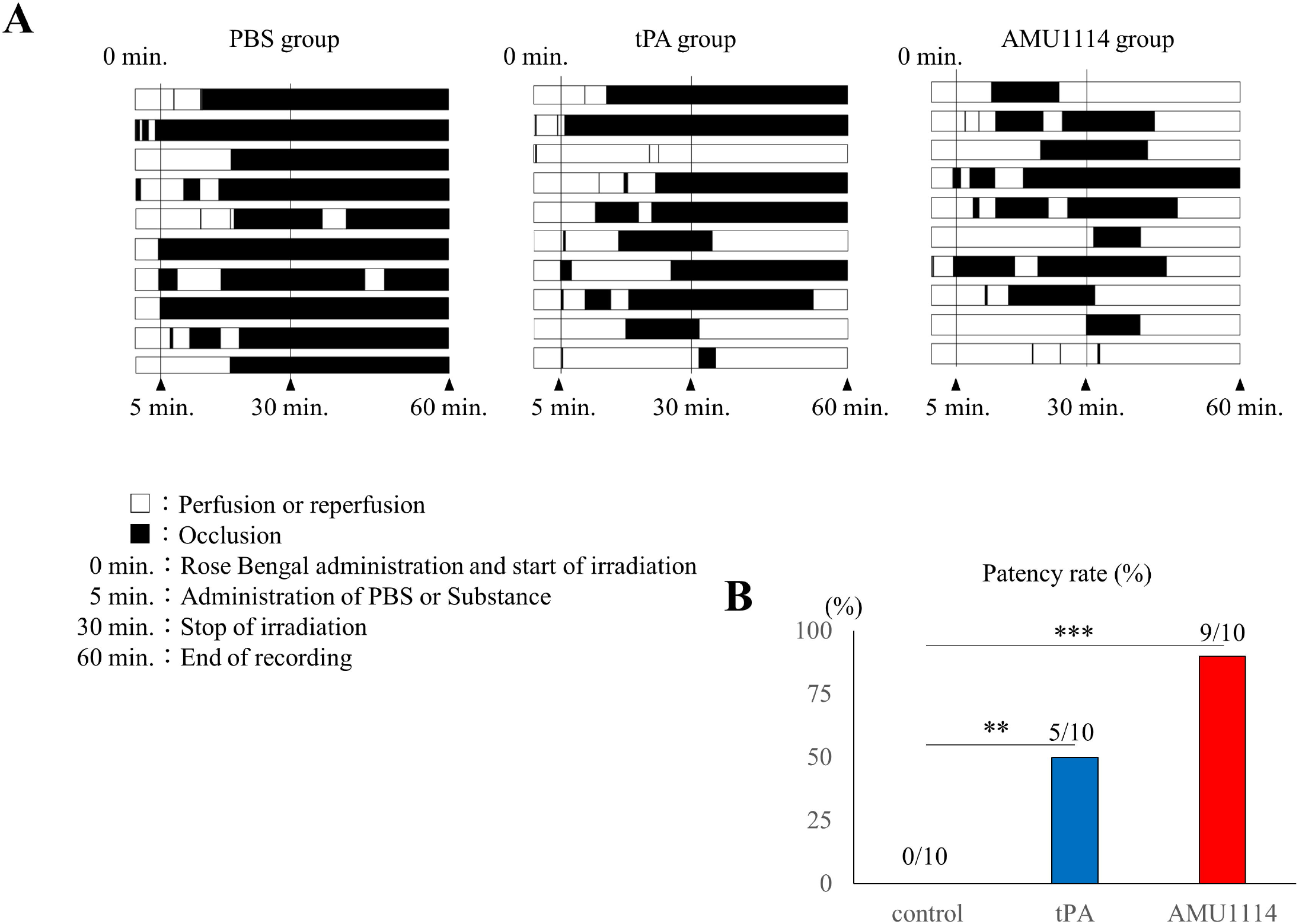
Effects of tPA and AMU1114 on mouse carotid artery embolism. (A) 0 minutes: Rose Bengal administration and start of irradiation; 5 minutes: administration of PBS, tPA, or AMU1114; 30 minutes: End of irradiation; 60 minutes: end of recording. ▪, occlusion; □ perfusion or reperfusion. (B) Number of mice (n = 10) with vascular patency at the end of measurement. Control vs tPA (*P* = .0098), control vs AMU1114 (*P* = .0001), and tPA vs AMU1114 (*P* = .051) (χ^2^ test).

## Discussion

In this study, we successfully constructed a fusion protein of mutated UK and anti-IF mAb, AMU1114, and confirmed that it retained the activity of UK. When UK and AMU1114 were administered to mice, the reduction in plasminogen was more strongly suppressed by AMU1114 than by UK, indicating that AMU1114 is safer than UK. The *in vitro* fibrinolytic activity of AMU1114 was almost the same as that of tPA. The bleeding toxicity was lower for AMU1114 than for tPA, according to the *in vivo* experiments of the effect of test substances to plasminogen consumption in blood of normal mice. In terms of the antithrombotic effect, a thrombus was formed in the carotid artery of mice by the PIT method, and the thrombolytic effect of the test substances was examined. Among the control, tPA, and AMU1114 groups, there was no significant difference in the time from the start of green light irradiation until complete occlusion of blood flow (TTO), although tPA and AMU1114 took longer than the control. Regarding vascular patency, tPA and AMU1114 had longer vascular patency times during blood flow measurements than the control. At the end of blood flow measurement, the number of individuals whose blood vessels were patent was 0/10 for controls, 5/10 for tPA, and 9/10 for AMU1114. Thus, tPA and AMU1114 had the effect of thawing the prepared thrombus and reopening blood vessels. At the same time, AMU1114 had a greater thrombolytic effect than tPA, as reflected by the larger number of individuals with reopened blood vessels. Our fusion protein has three main characteristics: first, because AMU1114 recognizes only IF, the mAb can be delivered to IF exclusively at lesion sites *in vivo* without being neutralized by precursors or degradation products in blood. Although the strategy of delivering plasminogen activators to the affected area with an antibody against anti-IF has been considered, our previous and present experiments confirmed that the antibody used for that purpose, 59D8, binds to IF and fibrinogen as well. In other words, the 59D8 antibody is neutralized by fibrinogen, which is much more abundant than IF *in vivo*, and may not be effectively delivered to the lesion. On the other hand, the antibody we have established, 1101, enables selective *in vivo* delivery by recognizing only IF; this is possible because the epitope of 1101 exists on the β-chain in the pit structure that is only present in IF. Second, AMU1114 acts selectively on IF in lesions. The plasminogen activator selectively converts plasminogen into active plasmin on IF.^23^ Plasmin is inactivated *in vivo* by endogenous inhibitors such as α2-PI, except when IF is present.^16^ Therefore, the UK moiety of AMU1114 can function selectively in lesions. Third, due to the mutation in the plasmin cleavage site, activated UK remains linked to AMU1114 on IF, and the fusion protein can function for a sufficiently long time in the lesion. Fortunately, this mutation does not decrease UK activity.

Thrombotic complications occur in patients with various infectious diseases and cancer.^1-3^ They may also be associated with worsening of clinical condition, e.g., fibrosis associated with elevated blood coagulation at local sites of infection and cancer. Although it may be difficult to use a thrombolytic agent in such situations, we believe that developing a safer formulation is a step in the right direction.

While additional data are accumulated, a research cell bank of this antibody-UK fusion protein should be established for the purpose of future clinical trials in patients with potentially fatal coagulopathy associated with cerebral infarction, myocardial infarction, cancers, and viral infection.

## Acknowledgments

This work was financially supported in part by National Cancer Center Research and Development Fund (26-A-14 and 29-A-9 to Y.M.); the Project for Cancer Research and Therapeutic Evolution from the Japan Agency for Medical Research and Development (AMED) (17cm0106415h0002 to Y.M.); the Takeda Science Foundation (to Y.M.); and the Kobayashi Foundation for Cancer Research (to Y.M.) We thank Dr. B Osman for technical support and Dr. G Ishii for advice on pathology. We also thank to Mrs. H Koike and M. Mizoguchi (Araake) for their significant contributions in producing the original anti-insoluble monoclonal antibody, Mrs. M Ohmaru for her technical assistance, and Mrs. H Shindo for her secretarial support.

## Author contributions

Y.M. developed the original concept for the study. S.H and Y.M. designed the experiments. S.S. and Y.M. conducted pharmacological studies, immunohistochemistry, and turbidity assays. S.H. and Y.M. discussed and wrote the manuscript. All authors reviewed and contributed to the final version of manuscript.

